# READv2: Advanced and user-friendly detection of biological relatedness in archaeogenomics

**DOI:** 10.1101/2024.01.23.576660

**Authors:** Erkin Alaçamlı, Thijessen Naidoo, Şevval Aktürk, Merve N. Güler, Igor Mapelli, Kıvılcım Başak Vural, Mehmet Somel, Helena Malmström, Torsten Günther

## Abstract

The possibility to obtain genome-wide ancient DNA data from multiple individuals has facilitated an unprecedented perspective into prehistoric societies. Studying biological relatedness in these groups requires tailored approaches for analyzing ancient DNA due to its low coverage, post-mortem damage, and potential ascertainment bias. Here we present READv2 (Relatedness Estimation from Ancient DNA version 2), an improved Python 3 re-implementation of the most widely used tool for this purpose. While providing increased portability and making the software future-proof, we are also able to show that READv2 (a) is orders of magnitude faster than its predecessor; (b) has increased power to detect pairs of relatives using optimized default parameters; and, when the number of overlapping SNPs is sufficient, (c) can differentiate between full-siblings and parent-offspring, and (d) can classify pairs of third-degree relatedness. We further use READv2 to analyze a large empirical dataset that has previously needed two separate tools to reconstruct complex pedigrees. We show that READv2 yields results and precision similar to the combined approach but is faster and simpler to run. READv2 will become a valuable part of the archaeogenomic toolkit in providing an efficient and user-friendly classification of biological relatedness from pseudohaploid ancient DNA data.

## Background

The analysis of biological relatedness has become an established part of the archaeogenomic toolkit [1,2]. It has provided us with important insights into the social structures of prehistoric groups [3–25], including Neandertals [26]. Furthermore, it serves as a quality control (QC) step in many bioinformatic pipelines to identify sample duplicates or exclude close relatives from population genomic analyses. This development has been facilitated by advances in both ancient DNA wet lab procedures and specifically designed bioinformatic methods, as the specific properties of ancient DNA do not allow the application of most approaches used with modern DNA [2]. Studies of biological relatedness in prehistoric groups are now reaching up to almost 100 individuals from the same site [22], highlighting the need for further development of methods to produce optimized and efficient tools in this area.

In 2018, we published READ (Relationship Estimation from Ancient DNA) [27] as one of the first tools specifically designed to infer biological relatedness from ultra-low coverage ancient DNA data. READ uses pseudohaploid input data and divides the genome into 1 Mbp windows, estimating the pairwise mismatch rate [28] per window and then using the genome-wide mean for relationship classification. The values are normalized by the expected pairwise mismatch rate for an unrelated pair of individuals from the same population to account for differences in background relatedness due to population diversity and SNP ascertainment. READ then uses this normalized pairwise mismatch rate (P0) to classify pairs of individuals as identical/twins, first-degree relatives (parent-offspring and full siblings), second-degree relatives (nephew/niece-uncle/aunt, grandparent-grandchild or half-siblings), or unrelated. This has been shown to work quite well with as little as 0.1x shotgun coverage per genome [27]. Recent years have seen the introduction or application of more advanced methods into the field which work with lower amounts of data (Table S1), provide resolution for more distant degrees of relatedness, and/or are able to differentiate between different types of relationships for the same degree (e.g. parent-offspring versus full siblings) [29–35].

Nevertheless, READ continues to be a popular tool in the field partly for its user-friendliness. Since READ uses pseudohaploid genotype calls as input, it allows the use of the same files used for other population genetic analysis without the need to generate files including sequencing read counts, calculate genotype likelihoods, or use imputation. Furthermore, READ has very simple assumptions estimating the expected pairwise mismatch rate from the data without the need for population allele frequencies, which allows using it as part of initial QC procedures or in populations (or species) for which little additional information is available.

READ [27] had been implemented as a Python 2 script, taking plain text Plink files (tped/tfam) as input. However, the last version of Python 2 was released in 2020 and some systems have already stopped supporting the language. Furthermore, READ wrote a large number of temporary files to the hard disk which were then analyzed by a separate R script called from the Python script. The output of the R script was then again read into Python and the final output was prepared. This back and forth between two scripts in different languages created ample possibilities for incompatibilities and unhandled errors. As READ continues to be used by many researchers in the archaeogenomics community, a reimplementation in Python 3 is warranted and provides the opportunity to add new features and improvements to its resource usage to be prepared for larger datasets.

Here, we re-implement the original READ [27] (READv1 hereafter) in Python 3 as READv2. The input file format has been changed from plain text Plink tped/tfam to binary Plink bed/bim/fam, requiring less space on the hard disk. All analyses are carried out within the same Python script using *NumPy* [36] and *pandas* [37] libraries, avoiding the excessive use of temporary files and the calling of a separate R script. Furthermore, by using a simulated dataset with known relationships and aDNA characteristics, we tested different window sizes which, in READv1, had been set to a default value without proper comparison. Consequently, we change the default values, obtaining a minor gain in accuracy. Finally, when the amount of data is sufficient, we add and test new features for classifying third-degree relatives and for distinguishing between siblings and parent-offspring when a pair has been classified as first-degree relatives.

## Results

### Resource demands

As some of the choices made during reimplementation are aimed at increasing the computational speed of READv2, we first test the resource demand using an empirical dataset. Rivollat et al. [22] recently reconstructed pedigrees from 94 individuals genotyped at ∼1.15 million autosomal SNPs. READv2 analyzed the 4371 pairs in this dataset in ∼8.5 minutes compared to nearly 5 hours for READv1 (Figure 1A). This substantial performance gain can be attributed to the use of binary input files, loading the full data into memory, and using *NumPy* [36] and *pandas* [37] for the analysis. The gain in running time comes at the cost of an increased memory demand (Figure 1B), but the required 5.64 GB is well within the standard resources provided by current personal computers. We also tested the resource demands for subsampled datasets. Subsampling to 50% of the individuals reduced the running time of READv2 to about one quarter (Figure 1A), suggesting a quadratic relationship due to the pairwise comparisons made. Subsampling 50% of the SNPs did reduce the running time by only about one-third. Subsampling had similar effects on memory usage even though the exact proportions of reduction differ slightly (Figure 1B). Using READv2 it was even feasible to analyze a full simulated dataset of 696 individuals and 241,860 pairs of individuals (see Methods and [38]) in less than 4 hours, but this required 237.5 GB of memory. This highlights that even for such extremely large datasets, READv2 provides an option to analyze the full dataset at once if enough memory is available (e.g. on clusters).

**Figure 1:**
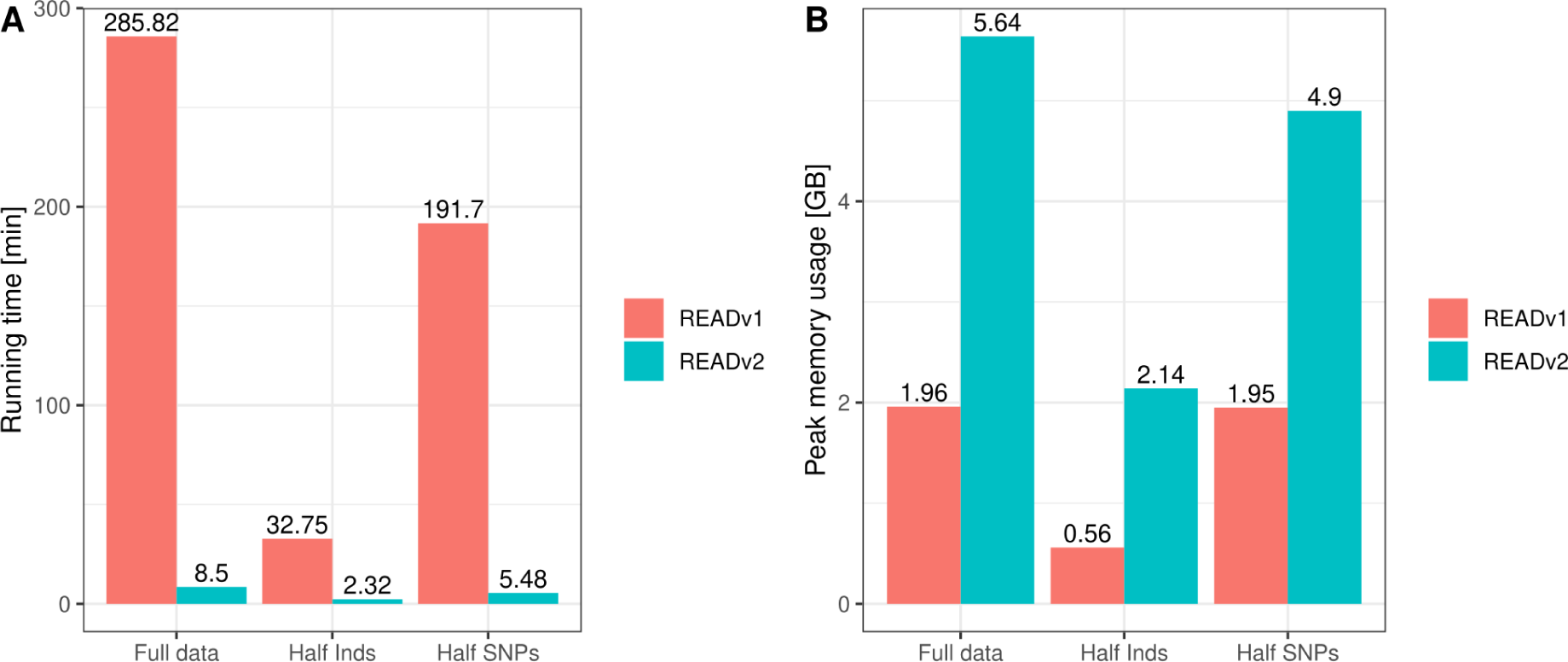
Time (A) and memory usage (B) comparison of READv1 and READv2 in a cluster node with two Intel Xeon E5 2630 v4 at 2.20 GHz/core CPUs and 128 GB RAM. The resource usage was tested with the dataset from Rivollat et al. [22] with 94 individuals. READv2 performs conspicuously better in terms of running time compared to READv1. However, READv1 uses slightly less memory for the same operations, because READv1 writes the results of intermediate steps to the hard drive and reads them back to perform further analyses instead of holding all the data in the memory as READv2 does.

### Window size

READv1 [27] used a default window size of 1,000,000bp which was inspired by GRAB [39], but it was never tested whether other window sizes could result in better accuracy. Therefore, we tested different window sizes (ranging from 100kbp to 20Mbp) on simulated data with known degrees of relationship [38]. In addition to this window-based approach where the test statistic P0 is estimated from the mean across all windows, we tested calculating a genome-wide P0 without splitting the genomes into separate windows. Interestingly, READv2 seemed to perform slightly better for smaller compared to larger window sizes, but overall the genome-wide estimate worked best (Figure 2, see Figure S1 for additional window sizes). The differences are more pronounced for second-degree relationships. At 0.05X and 0.1X, we observe high false positive rates for second- and third-degree relatedness as many unrelated pairs are classified into these categories (Figure S2). At 0.01X, unrelated individuals are even classified as first-degree or identical twins (Figure S2), resulting in a reduced false positive rate for second- and third-degree but an increased false positive rate for first-degree. Overall, READv2 performs well down to at least 0.1X sequence data in the simulated dataset. This corresponded to on average about 1,878 overlapping SNPs for each pair of individuals at an expected mismatch for unrelated individuals of ∼0.247 (Table 1). For the implementation of READv2, we set the genome-wide estimates as default, but users can adjust the settings if they wish to use different window sizes. All analyses below are based on the new default settings.

**Figure 2:**
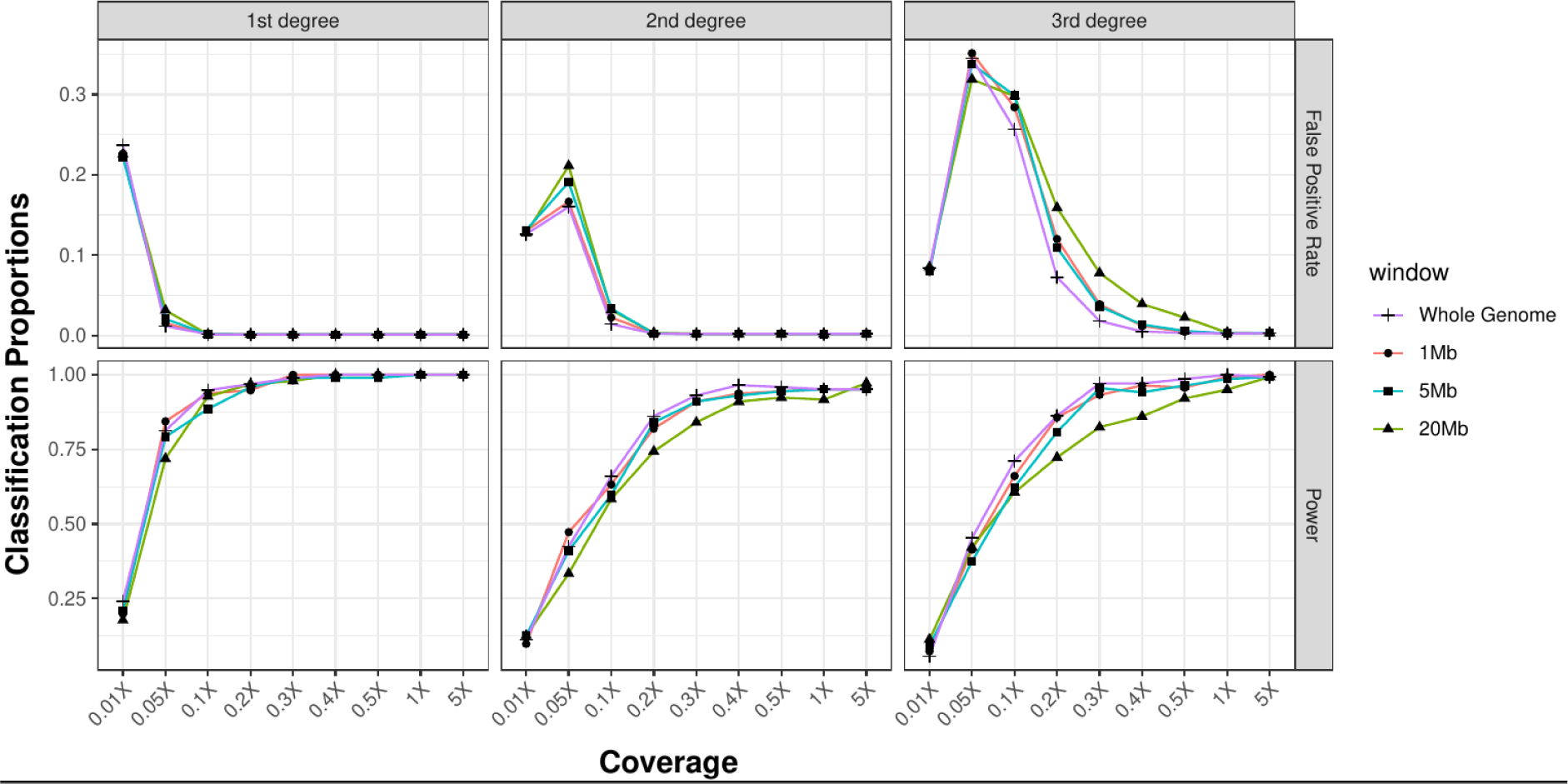
The power (i.e. the proportion of correctly classified pairs) and false positive rates (proportion of unrelated pairs classified into the respective degree) of READv2 assignment using simulated first-degree (n=118), second-degree (n=150), and third-degree pairs (n=144). The analyses were performed using varying window sizes (1Mb, 5Mb, 20Mb) (additional window sizes are shown in Figure S1) and for the genome-wide estimate (“Whole genome”), and also using varying coverages (0.01X, 0.05X, 0.1X, 0.2X, 0.3X, 0.4X, 0.5X, 1X, 5X). Classification proportions are shown in Figure S2. Overall, the genome-wide estimate performs better than any of the window-based methods.

**Table 1:** Number of overlapping SNPs in the simulated dataset (corresponding to the analysis shown in Figures 2 and S1)

| Simulated Coverage | Average number of overlapping SNPs [min, max] | Average number of effectively overlapping SNPs (Number of SNPs times expected pairwise mismatch of unrelated individuals) [min, max] |
| --- | --- | --- |
| 5X | 181,462 [174800, 181778] | 44,844 [43197, 44928] |
| 1X | 77,331 [57424, 78292] | 19,105 [14184, 19336] |
| 0.5X | 30,914 [21265, 31520] | 7,630 [5243, 7783] |
| 0.4X | 21,875 [14741, 22414] | 5,398 [3633, 5532] |
| 0.3X | 13,636 [8993, 14094] | 3,364 [2215, 3481] |
| 0.2X | 6,737 [4294, 7083] | 1,661 [1058, 1744] |
| 0.1X | 1,878 [1109, 2073] | 463 [273, 511] |
| 0.05X | 497 [275, 590] | 122 [68, 145] |
| 0.01X | 21 [6, 44] | 5 [1, 11] |

We need to note that the normalization value, i.e. the expected pairwise mismatch between unrelated individuals, can be seen as a useful approximation for the average amount of information per SNP in a dataset. The average mismatch for unrelated pairs is expected to reflect average heterozygosity under Hardy-Weinberg equilibrium for the SNP set used and it can vary substantially between populations and ascertainment schemes [27]. Therefore, an accurate assessment of the performance requires taking both the number of overlapping SNPs as well as the average mismatch for unrelated pairs into account. We use the product of these two values as the “effective number of overlapping SNPs” for our analyses below in order to make the number of SNPs needed more comparable across datasets with different SNP panels or population background diversities. The “effective number of overlapping SNPs” can thus be considered to represent a measure of the amount of information available for a pair to be used in kinship estimation.

### Third-degree relationship

For READv1, we did not introduce the option to classify pairs of individuals as third-degree relatives. One reason was that we expected most applications with very low coverage data, so the ranges for second-degree, third-degree, and unrelated pairs would overlap substantially, leading to false classifications. Furthermore, the 1000 Genomes Project [40] dataset that was used for testing only included a very limited number of third-degree relatives. Nevertheless, other researchers have modified READv1 to classify up to third-degree relatives [25,41], suggesting that the READ approach might be able to perform such classifications in certain situations. For Figure 2, we also tested the ability of READv2 to classify third-degree relatives. As expected, the third-degree classification requires more data than second- or first-degree classifications. For low amounts of data, we see third-degree relatives frequently being assigned to other categories and unrelated pairs being classified as third-degree (Figure S2). From about 0.3X sequencing coverage, power and false positive rates are similar to the values observed for first- and second-degree. Therefore, we decided to implement a threshold for the amount of overlapping data below which pairs falling into the third-degree category are automatically classified as “unrelated” while a third-degree classification is performed for larger amounts of data. Sequencing coverage of 0.3X corresponds to ∼13,600 overlapping SNPs in this simulated data or ∼3,400 “effectively overlapping SNPs” when this value is multiplied by the expected distance of unrelated pairs. To avoid false classifications in empirical data, we implement a conservative threshold of 5000 “effectively overlapping SNPs”, below which we do not attempt to classify third-degree relatives.

### Distinguishing between parent-offspring and siblings

Based on the estimate for a normalized pairwise mismatch rate, READ classifies pairs of individuals into degrees of relationship. For first-degree relatives, two options exist: parent-offspring and full siblings. Parent-offspring pairs share exactly one chromosome for each position of the genome while siblings should approximately share zero or two chromosomes for about one-quarter of the genome each, and one chromosome for the remaining half of the genome. By plotting the variation in the pairwise mismatch rate across the genome, some studies have resolved individual pairs of first-degree relatives [22,42], while ancIBD [35] and KIN [32] implicitly model this as part of their Hidden Markov Models (HMM). We explored whether READv2 could use the empirical distribution across windows to distinguish between parent-offspring and full-sibling relationships. Larger windows appear more suitable for this purpose (Figures S3 and S4), so we perform this analysis with 20Mb windows. In default settings, READv2 will assess the degree of relationship based on a genome-wide estimate of the pairwise mismatch rate, followed by a separate round of classification for first-degree relatives based on 20Mb windows. As a test statistic, we use the proportion of windows classified as unrelated (i.e. no shared chromosome) or identical (i.e. both chromosomes shared), corresponding to Cotterman coefficients k0 and k2, respectively. As expected, this proportion is low for parent-offspring and around 0.5 for full siblings when sufficient data are available (Figure 3). For low amounts of data, the proportion of k0 and k2 windows first starts to increase for parent-offspring pairs and later also for siblings. While the two types are well separated down to 0.5X coverage in the simulated dataset (or ∼8,000 “effectively overlapping SNPs”), they overlap at 0.2X and below. We used these results to set thresholds for the separation of parent-offspring from full siblings based on the proportion of windows classified as unrelated or identical. A pair of first-degree relatives is classified as parent-offspring if the proportion is below 0.3, as siblings if the proportion is between 0.35 and 0.6, and as “N/A” otherwise. Since low amounts of overlapping data result in proportions >0.6, this allows us to avoid a classification if the amount of data is insufficient. There is, however, the risk that parent-offspring would be classified as full siblings for a specific range of overlapping data (between ∼1600 and ∼5000 effectively overlapping SNPs).

**Figure 3:**
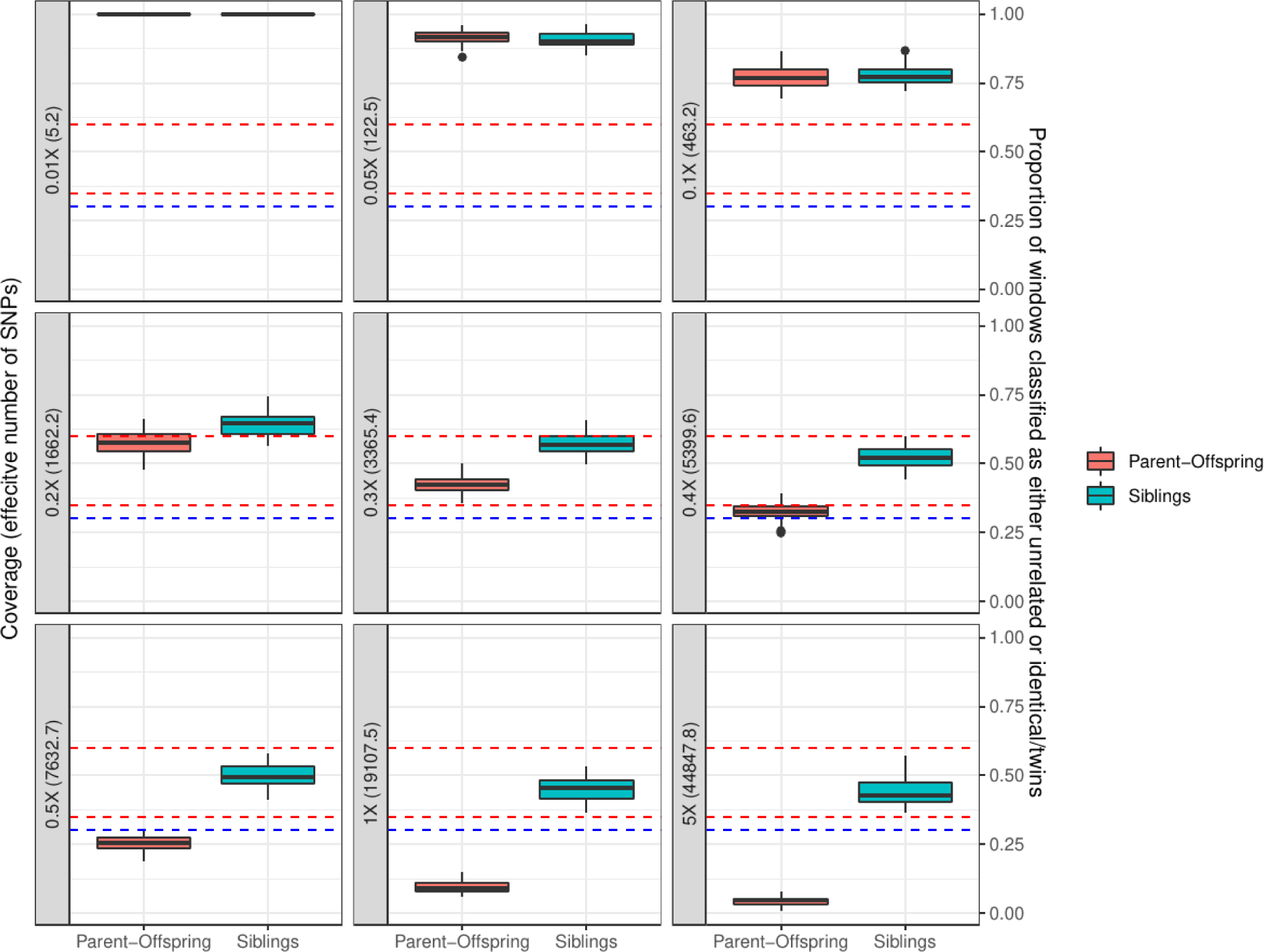
Proportion of windows that are classified as either unrelated or identical/twins. The analysis was done by using a window size of 20Mb with 68 parent-offspring and 49 sibling pairs. Dashed lines indicate the thresholds chosen to distinguish between parent-offspring and siblings in the classification. The area under the blue dashed line shows the “parent-offspring” zone, while the area between the red lines presents the “siblings” zone. The separation is clear for coverages over 0.5X and roughly 8,000 effectively overlapping SNPs. As the coverage and the number of effectively overlapping SNPs reduce, the distributions begin to overlap and the proportions increase overall. Note that the average effective number of overlapping SNPs is slightly different from Table 1 as a different subsampling of the full dataset was used for the analysis in Figure 2 and Table 1.

To test this feature in an independent dataset, we selected SNP genotype data from the 1000 Genomes Project where individuals from different populations have been genotyped using the Illumina Omni2.5M chip HD genotype SNP array including 2,458,861 SNPs. We selected the populations CHS (Han Chinese South) and YRI (Yoruba) which contained the largest number of first-degree relatives: 105 parent-offspring pairs and 8 sibling pairs for CHS, and 112 parent-offspring pairs and 4 sibling pairs for YRI. The overall number of full siblings in the dataset is low, not allowing for proper testing of the feature. However, as siblings would be classified as “N/A” for increased noise, the critical test is whether parent-offspring pairs are classified as siblings at reduced amounts of data. These tests have been performed in each population separately. Similar to the simulations, parent-offspring pairs are correctly classified when the amount of overlapping data is large (>14,000 effectively overlapping SNPs, Figure 4). Below this point, initially, parent-offspring pairs are increasingly classified as “N/A”. From 8,000 effectively overlapping SNPs and less, however, we see substantial numbers of parent-offspring pairs classified as siblings. For very low amounts of overlapping data (< 2,000 effectively overlapping SNPs), all are classified as “N/A”. These results are very similar to the results seen for the simulated data. To be conservative and avoid wrongly classifying parent-offspring pairs as siblings, we implement a default cutoff of 10,000 effectively overlapping SNPs below which classification is not performed.

**Figure 4:**
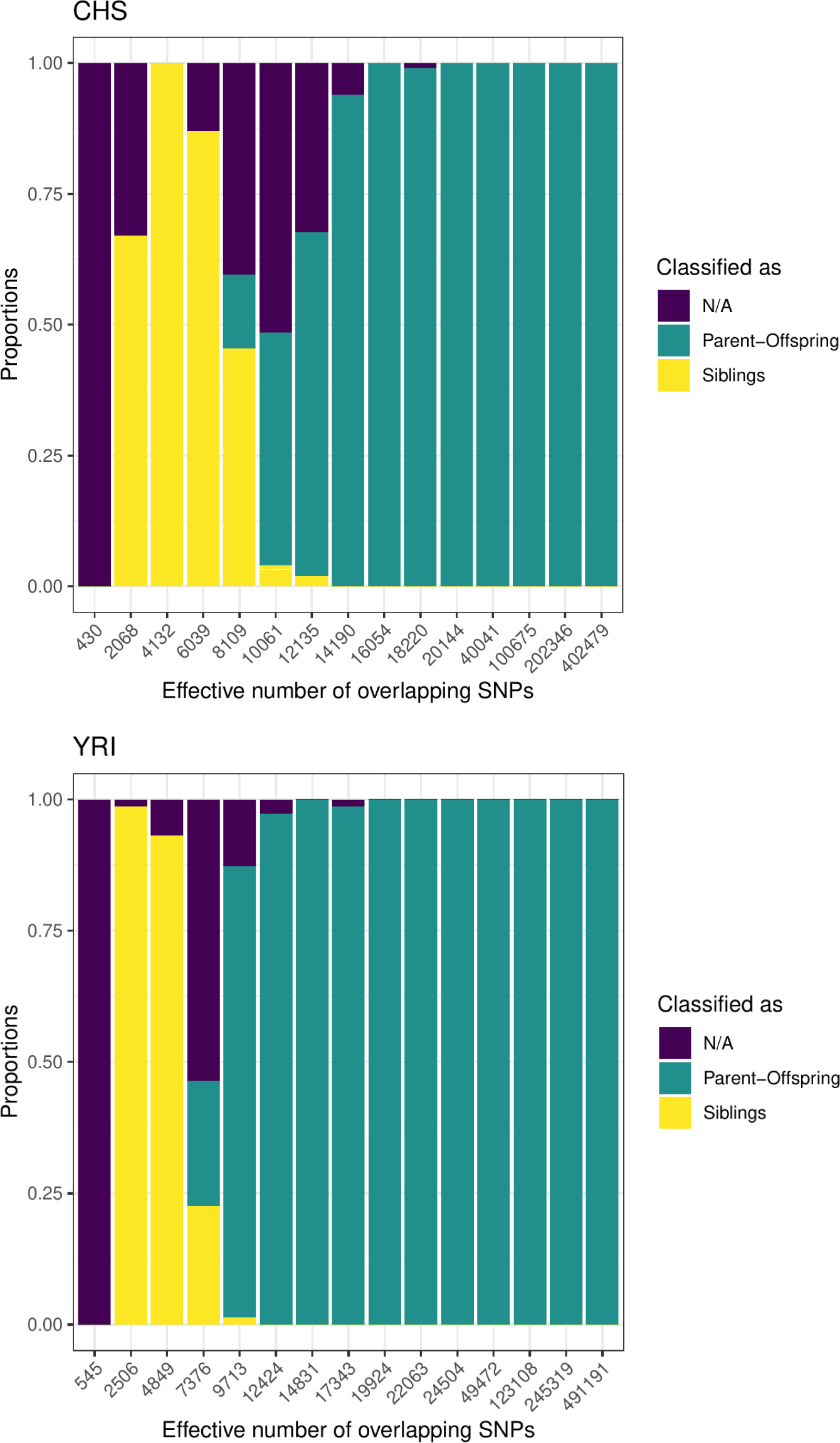
Classification of known parent-offspring pairs in empirical data. The feature was tested with CHS (Han Chinese South) and YRI (Yoruban) populations from the 1000 Genomes Project for different amounts of overlapping SNPs (n=105 and 122, respectively). Similar to the result of the analysis made with the simulated data, parent-offspring pairs are correctly classified for high amounts effectively overlapping SNPs for both populations. As this number reduces, more siblings and N/A classifications start to be seen.

### Empirical data application: Rivollat et al. (2023)

A recent study conducted the tremendous effort of obtaining genome-wide data for 94 individuals from the same site in Neolithic France, Gurgy ‘les Noisats, and then reconstructing pedigrees for the individuals buried at the site [22]. The authors first ran READ [27] to estimate the degree of relatedness for each pair of individuals, followed by lcMLkin [29] to differentiate between parent-offspring and siblings among first-degree relationships. They further used BREADR [33] for individual pairs as well as imputation and ancIBD [35] for higher degrees. We have already shown that READv2 can process this dataset a lot faster than READv1 (Figure 1A). By adding the new feature to differentiate parent-offspring and siblings, we are able to perform the analysis that originally needed two different tools with different input files in a single analysis that is orders of magnitude faster than the first step of the original analysis alone. About 94% of the pairs of individuals in this dataset have more than 40,000 overlapping SNPs at an expected pairwise mismatch rate for unrelated individuals of 0.245, i.e. the product for most of them is >10,000 representing a situation where the new feature of READv2 should be applicable. All 86 first-degree pairs identified by READv1 in the original study were confirmed by READv2 (Supplementary Data 1). For 81 of them, READv2 was able to discern parent-offspring and siblings, all in agreement with the pedigree in [22]. Four of the remaining five were not classified due to their low amounts of overlapping data (less than 7,000 effectively overlapping SNPs), and of these, two parent-offspring pairs would have been classified as siblings if the threshold had not been in place. Further, due to low coverage, two of these four were only classified by context in [22] rather than by a clear signal in the classification softwares. The fifth pair had sufficient data but fell between the ranges of sibling and parent-offspring used by READv2.

Notably, both READv1 and READv2 identified one additional pair of first-degree relatives (GLN207A-GLN279) that the original study did not detect with READv1. This likely reflects the stochasticity of random sequencing read sampling in the independent genotype calls as the original study had them just above the first-degree classification threshold, while our results have them just below the threshold. Rivollat et al. had them as siblings in their pedigree based on the classification of other relatives and the lcMLkin results. READv2 also classified them as siblings. Another notable pair is GLN285A-GLN285B, which lcMLkin had as an outlier suggestive of a sibling relationship. READv2 classified them as parent-offspring. Rivollat et al. did not directly classify them as parent-offspring but excluded a sibling relationship due to the presence and absence of relationships with other individuals. Another pair, GLN288-GLN289B, was not classified by lcMLkin due to the low coverage of GLN289B. READv2 classified this pair as parent-offspring as also concluded by [22] due to the classification of other related individuals.

This re-analysis of the data from Gurgy ‘les Noisats’ [22] illustrates that READv2 alone can lead to very similar results as the combination of READv1 and lcMLkin in the original study. Both of these latter approaches appear to miss some cases that were only resolved through context or by excluding certain types of relationships with additional data (e.g. uniparental markers, age at death). Both approaches also fail in similar cases due to low amounts of overlapping data. This highlights that READv2 can be used in such studies to resolve large pedigrees when combined with additional data. READv2 has the advantage that it is much faster than the combined approach and all results can be obtained by running a single tool.

## Discussion

We introduce a new version of the popular tool for inferring biological relatedness from ancient DNA data, READ. Firstly, READv2 can be described as a Python 3 re-implementation of READv1 with substantially improved running times. The implementation in a single language should also increase portability and avoid possible version conflicts. Secondly, READv2 has an updated default behavior as the pairwise mismatch rate is not derived from the mean across genomic windows but as a genome-wide estimate, leading to up to 5% improvement in classification accuracies. Finally, we added two new features: the ability to classify up to third-degree relatives, which requires at least 5000 effectively overlapping SNPs, and the ability to differentiate between different types of first-degree relationships, *viz.* full siblings and parent-offspring, which requires at least 10,000 effectively overlapping SNPs. The introduction of “effectively overlapping SNPs” as a measurement of the amount of available data should also make studies and datasets more comparable and increase the possibility of generalizing from benchmarking results as previous studies mostly compared the raw number of overlapping SNPs or sequencing depth without taking the information content per SNP into account.

We introduced READv2 as a version with increased efficiency and compared it to READv1. Other studies have already covered comparing the general READ approach to other methods used in the field [31–33,43,38]. Methods such as ancIBD [35], KIN [32], TKGWV2 [31] as well as the genotype likelihood-based lcMLkin [29] and ngsRelate [30] have specific advantages, either providing more precise results with lower amounts of data or by being able to detect higher than second-degree relatedness confidently. In contrast to READ, they often require additional data preparation and/or information, such as read counts, estimation of genotype likelihoods, imputation, or population allele frequencies, which are often difficult to obtain for aDNA data or simply not available for certain populations. ancIBD [35] and KIN [32] were both specifically designed for ancient DNA data and their HMM approaches allow for the classification of higher degrees of relatedness as well as the differentiation between siblings and parent-offspring pairs. READv2 is very similar in its approach to BREADR [33] and TKGWV2 [31], with each tool having its own unique feature. READv2 has the functionality to separate the different first-degree relationships, BREADR has a better quantification of uncertainty, and TKGWV2 works well with lower amounts of input data. We expect READv2 to find its own niche in this ecosystem of different methods. The combination of increased efficiency and READv1’s user-friendliness qualify it as a QC step in data processing pipelines or as the first tool in an analysis of biological relatedness, which can be followed up with other tools to detect more fine-scale patterns or to verify results. Adding the possibility of differentiation between siblings and parent-offspring pairs when sufficient amounts of data are available provides additional value for such an initial analysis. This feature requires about 10,000 “effectively overlapping SNPs”, which, assuming the popular 1240K SNP capture panel with 1.15 million autosomal SNPs and European Neolithic populations, would correspond to about 0.2X coverage per individual which is the case for a large proportion of all published human ancient genome-wide data [44]. Furthermore, the increased resolution of potentially classifying individuals as third-degree relatives for larger amounts of overlapping data (>5000 effectively overlapping SNPs) will improve the reconstruction of more complex pedigrees. Finally, we expect that the substantially improved running times make READ analysis feasible for future data sets, which will undoubtedly increase in sample sizes.

## Material and Methods

### READ Re-implementation

READv1 [27] was written in Python2 and R, with an R script called from the Python script to carry out specific analyses. A description of the READ workflow can be found in the Introduction section. The first step of this project was to re-implement READv1 in Python3, in order to update the script, increase efficiency and portability, and avoid possible version conflicts. The R script parts of READv1 were implemented using the Pandas library [37] in Python 3. Furthermore, with the reimplementation, the input file format was changed to binary PLINK bed/bim/fam files using the *PLINKIO* library (https://github.com/mfranberg/libplinkio). In order to avoid excessive loops and improve the method’s runtime, the pairwise comparison was implemented with the NumPy [36] library. In addition to the window-based approach for estimating the pairwise mismatch rate (P0), a single genome-wide estimate using all covered sites was implemented. In this case, the uncertainty for the pairwise mismatch rate is estimated using a block-jackknife approach with block sizes of 5 Mb as commonly employed in human population genomic studies [45].

For READv1, the classification thresholds were set to the mid-point between the expected P0 values for each degree. Consequently, we also set the cutoff for third-degree classifications halfway between the expected P0 for unrelated individuals (i.e. 1.0) and third-degree relatives (0.9375). Pairs of individuals with a normalized P0 between 0.90625 and 0.96875 are now classified as third-degree relatives if the number of effectively overlapping SNPs (number of overlapping SNPs times the pairwise mismatch rate expected for unrelated individuals) is 5000 or higher, otherwise they are classified as unrelated.

To differentiate between parent-offspring and sibling pairs, the genome is divided into windows of 20Mb and the classification is made based on the proportion of windows that are classified as either “identical/twin”, “unrelated”, or “third degree”. If that proportion is less than 0.3, the pair is classified as “parent-offspring”; if it is between 0.35 and 0.6, the pair is classified as “siblings”. For other proportions, or when the number of effectively overlapping SNPs is below 10,000, the type is not specified beyond “first-degree”. Furthermore, we expanded the output table by including information such as the number of overlapping SNPs, the number of effectively overlapping SNPs, and the kinship coefficient *θ* (= 1 - normalized P0) to fulfill several user requests. An overview of the READv2 workflow can be found in Figure 5. READv2 is available with instructions on its usage at: https://github.com/GuntherLab/READv2

**Figure 5:**
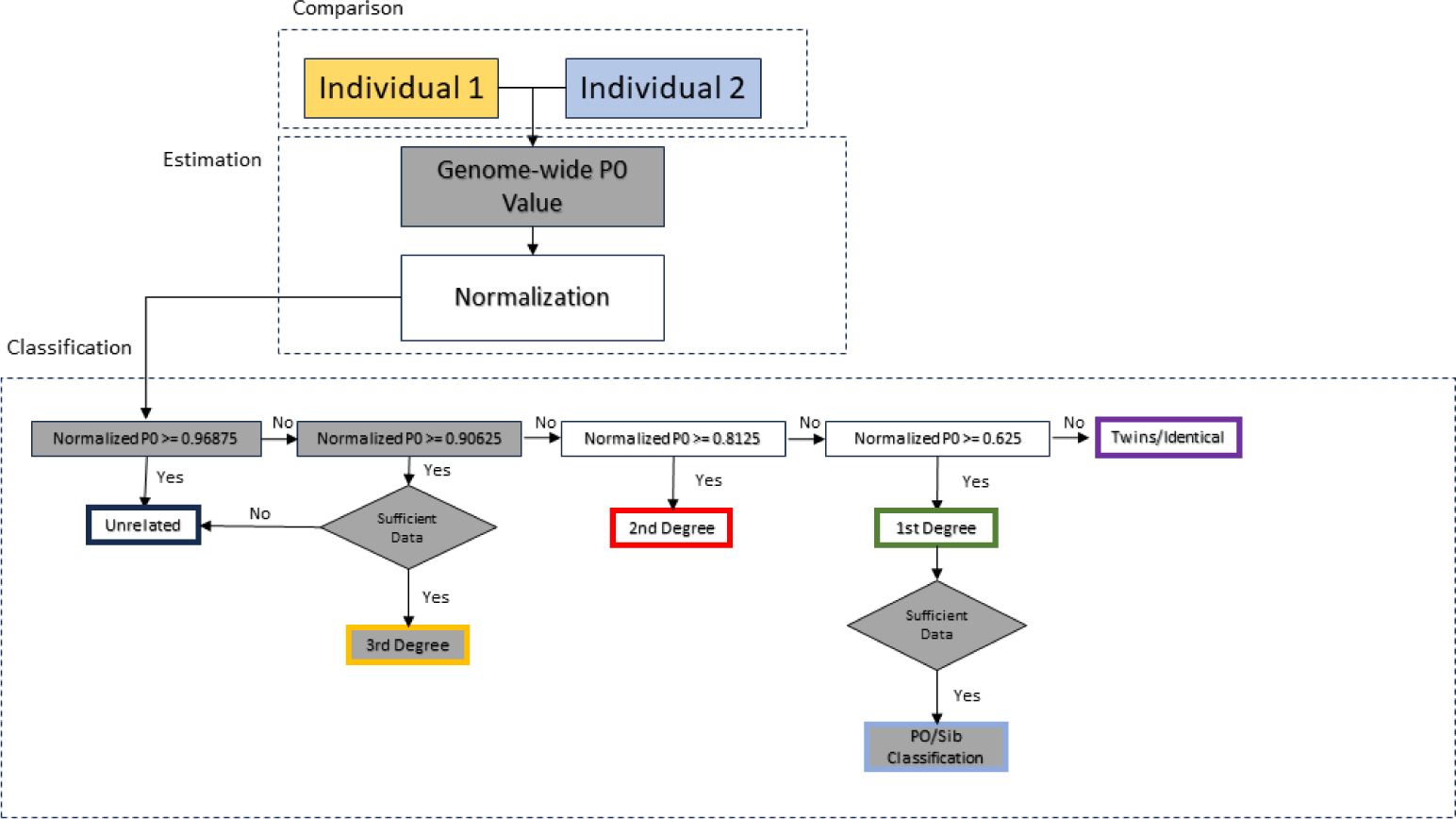
Flowchart of READv2. The novel steps and classification results that differ from READv1 have been highlighted in grey.

### Simulated pedigree data

The next step after the reimplementation was to create a benchmark with simulated data with known relationships to test the performance of READ. We used simulated ancient genome data representing pairs of individuals with known relationships [38]. Specifically, we applied simulation software *PED-SIM* (v 1.3) [46] to produce genotypes from pedigrees of various relationship degrees and types separately, including first-, second-, and third-degree relatedness. We created founder genotype data from scratch as follows: We chose SNPs with minor allele frequencies (MAF) equal to or higher than 0.01 of the modern-day Tuscany (TSI) sample (n=112) from the WGS data of the 1000 Genomes Project [47], and used them for the founder data generation. For the resulting 8,677,101 biallelic autosomal SNPs, we recorded the reference or alternative alleles at each SNP position as observed in the TSI dataset and calculated the alternative allele frequency (AAF) per SNP in TSI. We then created the genotypes of each founder by randomly choosing, for each SNP independently, the alternative or reference allele with probability AAF and 1-AAF, respectively, and repeating this twice to create a diploid genotype per founder. Note that this method of creating diploid genotypes eliminates any background relatedness among founders as well as any homozygosity tracts within founder genomes. We repeated the creation of founder data 12 times (or runs), each producing different sets of founders.

We independently generated 120 unrelated founders (10 individuals per run for n=12 runs) used for first-degree and 240 unrelated founders (20 individuals per run for n=12 runs) for second- and third-degree pedigree simulations. A sex-specific genetic map [48] with linear interpolation and crossover interference model [49] was used to simulate genotype data of related individuals using the “*--m*” and “*--intf*” options, respectively. We provided all possible sex combinations for the relationships in the *PED-SIM* parameter file (def file) using the “*--d*” option. In this way, we simulated distinct pedigrees with the same structure but varied with respect to the sex of the individuals. Additionally, we assigned the sex of the founders with the “*--sexes*” option. We also used the “*--keep_phase*” parameter to keep phase information in the output, the “*--miss_rate 0*” parameter not to introduce missingness, and the “*--founder_ids --fam*” parameters.

We simulated 72 pedigrees for first-degree relationships, 96 for second-degree relationships, and 96 for third-degree relationships. The founders of each pedigree and simulated individuals from distinct pedigrees were treated as “unrelated”. For each relationship type, we chose n=48 pairs. For instance we simulated n=72 individuals (n=24 trios) for parent-offspring relationships, resulting in 48 unique pairs. Consequently, the dataset comprises 696 individuals (n=72 for parent-offspring, grandparent-grandchild, and great-grandparent-great-grandchild and n=96 for siblings, half-sibling, avuncular, first cousin, and grand avuncular relationships).

### Simulated ancient DNA sequencing and processing

We used the *gargammel* software [50] to simulate ancient DNA-like read data. This ancient read simulator cuts a given FASTA file into short sequences of variable lengths, which reflect the distribution of actual ancient read lengths, and adds post-mortem DNA damage. Then, it adds Illumina adapters to the ends of the reads. Finally, sequencing errors and quality scores are introduced, producing ancient-looking FASTQ files. To generate input FASTA files for *gargammel*, for each individual separately (two files for each individual), at each SNP position, we inserted alternative alleles according to their genotype into the human reference genome (GRCh37) via *vcftools consensus* [51]. We then cut the FASTA files into 100 bp sequence intervals surrounding each SNP (50 bp on each side) using *bedtools getfasta* [52]. This step was performed because the most time-consuming step of the dataset preparation is mapping a vast number of genomes to the reference genome; limiting the number of reads to those surrounding the variable sites allowed significantly reducing the computation time. For aDNA read size distribution, we used the size distribution file (*sizedist.size*) from *gargammel*, but we removed values higher than 120 bp, resulting in a distribution with a mean of 66.2 bp and a median of 61 bp, ranging between 35 bps and 119 bps. Using Briggs model parameters, we specified the deamination patterns (-damage 0.024, 0.36, 0.009, 0.55) [53]. We set the final depth of coverage of the samples as 5x without any present-day human or microbial contamination.

We processed the *gargammel*-simulated read data following the same procedure as applied to ancient genome sequencing in the field [54]. Firstly, we removed the adapters from the simulated ancient reads and then merged the paired-end reads only if they had at least 11 base pairs (bp) length overlap using *AdapterRemoval v2.3.1* [55]. Secondly, the generated single-end ancient reads were mapped to a human reference genome (hs37d5) using the *bwa* software (v0.7.15) [56] with the “*aln*” option, and the non-default parameters “*-l 16500*”, “*-n 0.01*” and “*-o 2*”. We eliminated the reads with a minimum of 10% mismatches to the human reference genome. Finally, the remaining reads were trimmed ten bps from both ends to remove C-to-T substitutions (or reciprocally G-to-A) resulting from post-mortem damage of aDNA using the *bamUtil* software with the “*trimBAM*” option [57]. BAM files are available from Zenodo (https://zenodo.org/doi/10.5281/zenodo.10079684 and https://zenodo.org/doi/10.5281/zenodo.10079624) [38]. Genotypes were called from the BAM files using ANGSD v0.933 [58] with the options −checkBamHeaders 0 −doHaploCall 1 −doCounts 1 −doGeno ™4 −doPost 2 −doPlink 2 −minMapQ 30 −minQ 30 −doMajorMinor 1 −GL 1 −domaf 1. Pseudohaploid tped/tfam files were then generated with the ANGSD tool haploToPlink and converted to binary Plink files using Plink [59].

### Benchmarking

In order to reduce the memory and runtime of the window size comparisons, the dataset was divided into groups of 70 by involving all related individuals, i.e. all 3 individuals (two parents and one offspring) in a parent-offspring relationship, in a group with Plink –keep-fam command. The normalization value was calculated as the median mismatch per subsample of the data. To test the performance of READ for different coverages, the original simulation data was downsampled with SAMTOOLS view −s [60]. In order to see how window size affects the results and compare the window-based and genome-wide approaches, the power of the method (TP/(TP+FN)) and the proportion of unrelated pairs classified as related (false positive rate) were calculated for each coverage and window size (results shown in Figure 1).

While the window size comparisons were performed on the dataset separated into groups of 70, the tests for distinguishing between siblings and parent-offspring were conducted on the full dataset at once.

Empirical data from The 1000 Genomes Project [40] with known relationships have been used for further testing. The autosomal Illumina Omni2.5M chip HD genotype SNP array data consists of 2,368 individuals from 15 different populations with 2,458,861 SNPs. The populations with the most parent-offspring pairs, namely YRI (Yoruba in Ibadan, Nigeria) and CHS (Southern Han Chinese, China), were selected for further steps. The populations were separated into different .bed files with the PLINK –keep-fam option and later SNPs were down-sampled with the PLINK –thin option. Since the data was from modern samples and contained diploid genotype calls, the data were made homozygous by randomly selecting one allele at each position.

### Empirical data application

We downloaded 1240K SNP capture BAM files for 94 individuals excavated in Gurgy ‘les Noisats’ [22,61] from the European Nucleotide Archive [62]. Genotypes at ∼1.15 million autosomal SNPs were called using ANGSD v0.933 [58] with the options −checkBamHeaders 0 −doHaploCall 1 −doCounts 1 −doGeno ™4 −doPost 2 −doPlink 2 −minMapQ 30 −minQ 30 −doMajorMinor 1 −GL 1 −domaf 1. Pseudohaploid tped/tfam files were then generated with the ANGSD tool haploToPlink which were converted to bed/bim/fam with Plink v1.90b4.9 [63]. We then ran READv2 in default settings. To compare the resources needed from running READv1 and READv2, we also halved the number of individuals (with the Plink command –thin-indiv 0.5) and the number of SNPs (with the Plink command –thin 0.5).

## Acknowledgements

We thank members of the Human Evolution program, especially the Malmström and Günther labs, at Uppsala University for fruitful discussion, and especially Tiina Mattila and Jules Koelman for testing. We also want to thank Toomas Kivisild and Iñigo Olalde for encouraging us to explore third-degree relationships. HM and TG were funded through a Riksbankens Jubileumsfond grant (P21-0266). The computations and data handling were enabled by resources in projects SNIC 2022/22-1033, SNIC 2022/23-543 and NAISS 2023/22-1182 provided by the National Academic Infrastructure for Supercomputing in Sweden (NAISS) and the Swedish National Infrastructure for Computing (SNIC) at Uppmax, partially funded by the Swedish Research Council through grant agreements no. 2022-06725 and no. 2018-05973. Work on the simulated pedigree was performed through support by the European Research Council (ERC) Consolidator grant “NEOGENE” (no.:772390).

## Supplementary Figures

**Figure S1:**
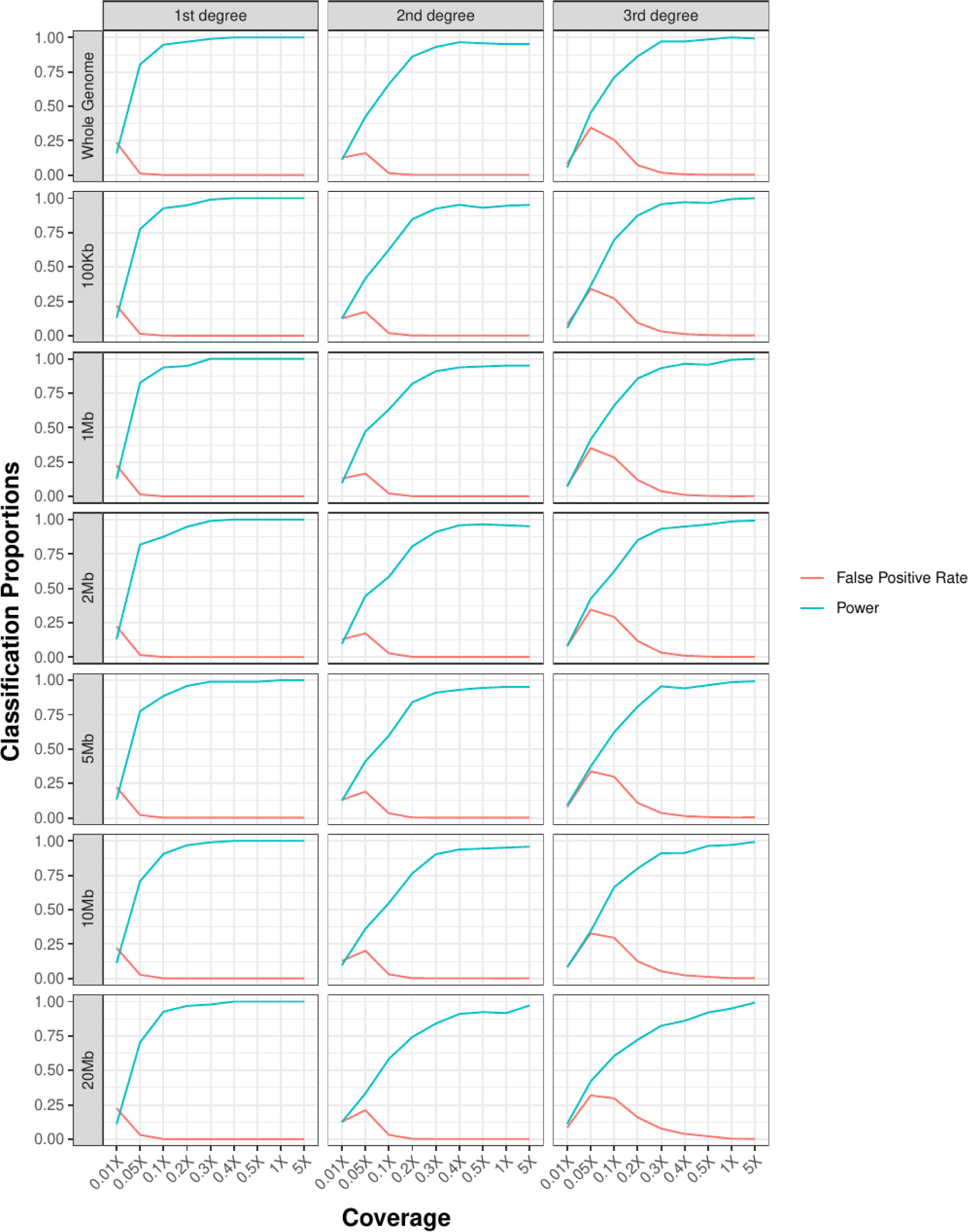
The power and false positive rates of READv2 for first-degree, second-degree, and third-degree pairs with additional window sizes. As also shown in Figure 2, READ performs well for coverages over 0.1X to classify first-degree pairs and over 0.3X for second- and third-degree pairs. Although there is not much difference between window sizes, the genome-wide estimate works the best overall.

**Figure S2:**
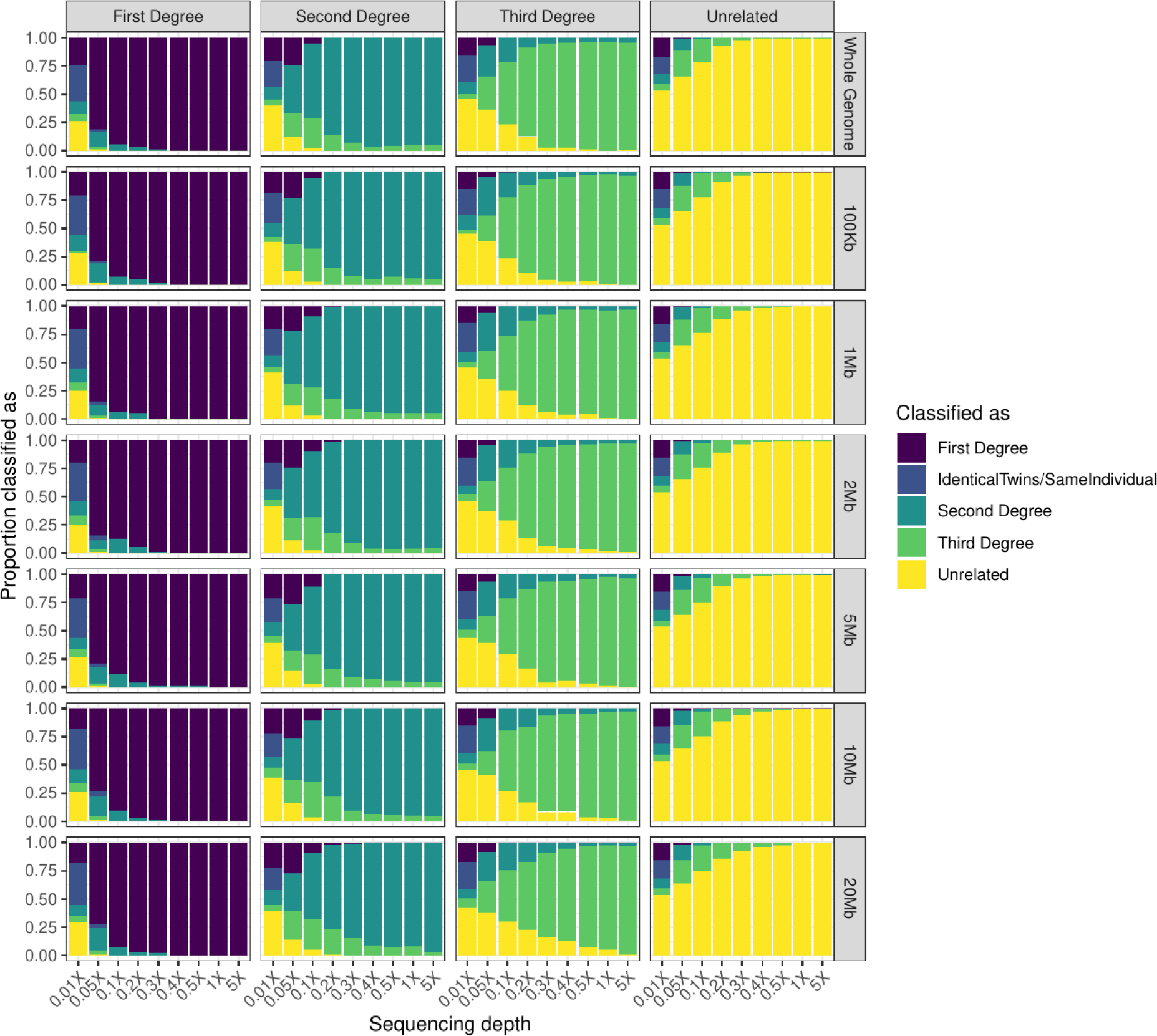
Proportions of simulated individuals with known biological relatedness classified into the different categories. Columns correspond to the true relationship while rows show the different window sizes, and colors represent the classification outcomes.

**Figure S3:**
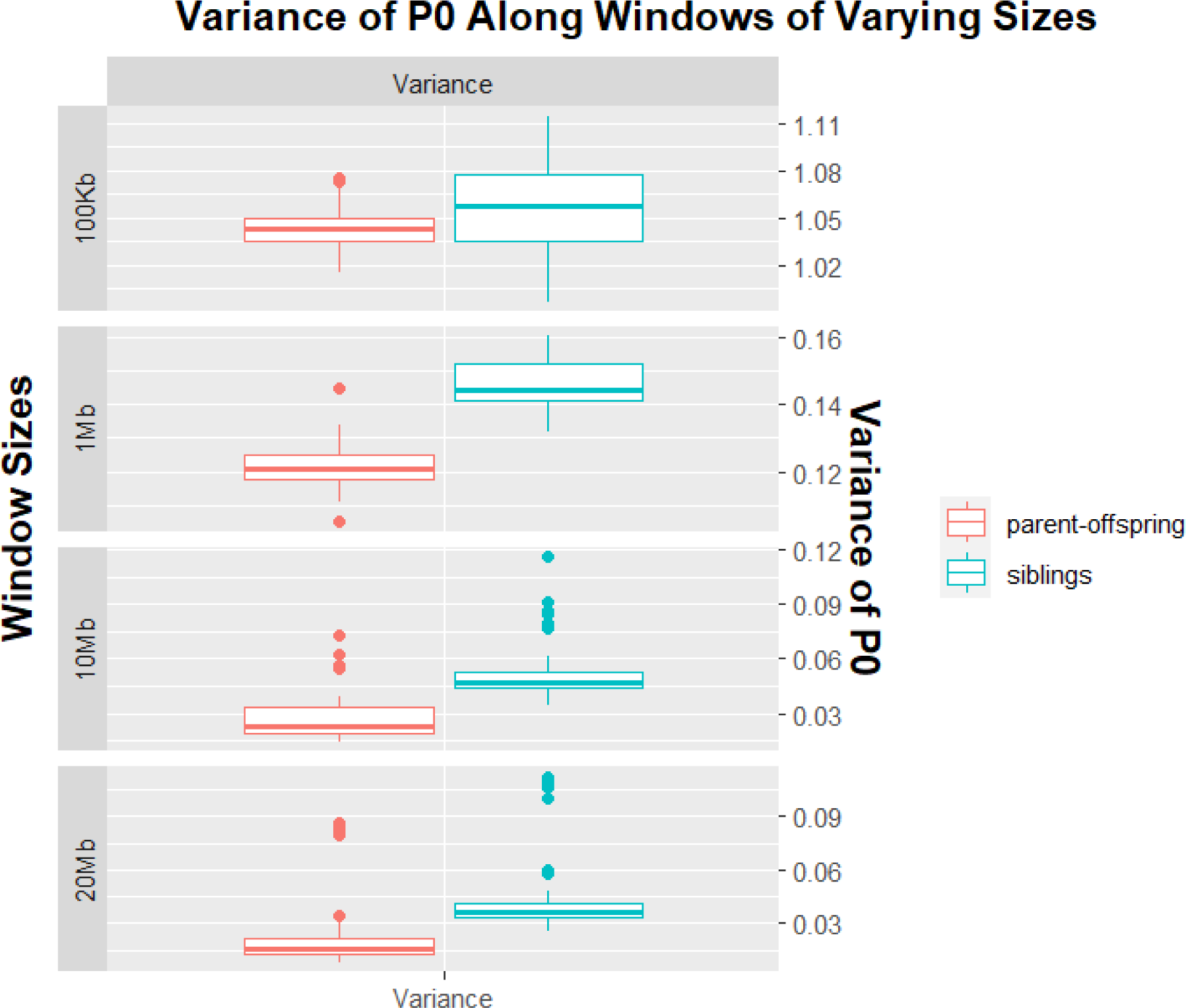
Variance of normalized P0 values along the windows of varying sizes for 1X coverage. The variance of parent-offspring and sibling pairs are visibly separated for large window sizes (1Mb, 10Mb, and 20Mb). However, as the window size decreases, that clear separation is lost. Moreover, the scale differs between window sizes used.

**Figure S4:**
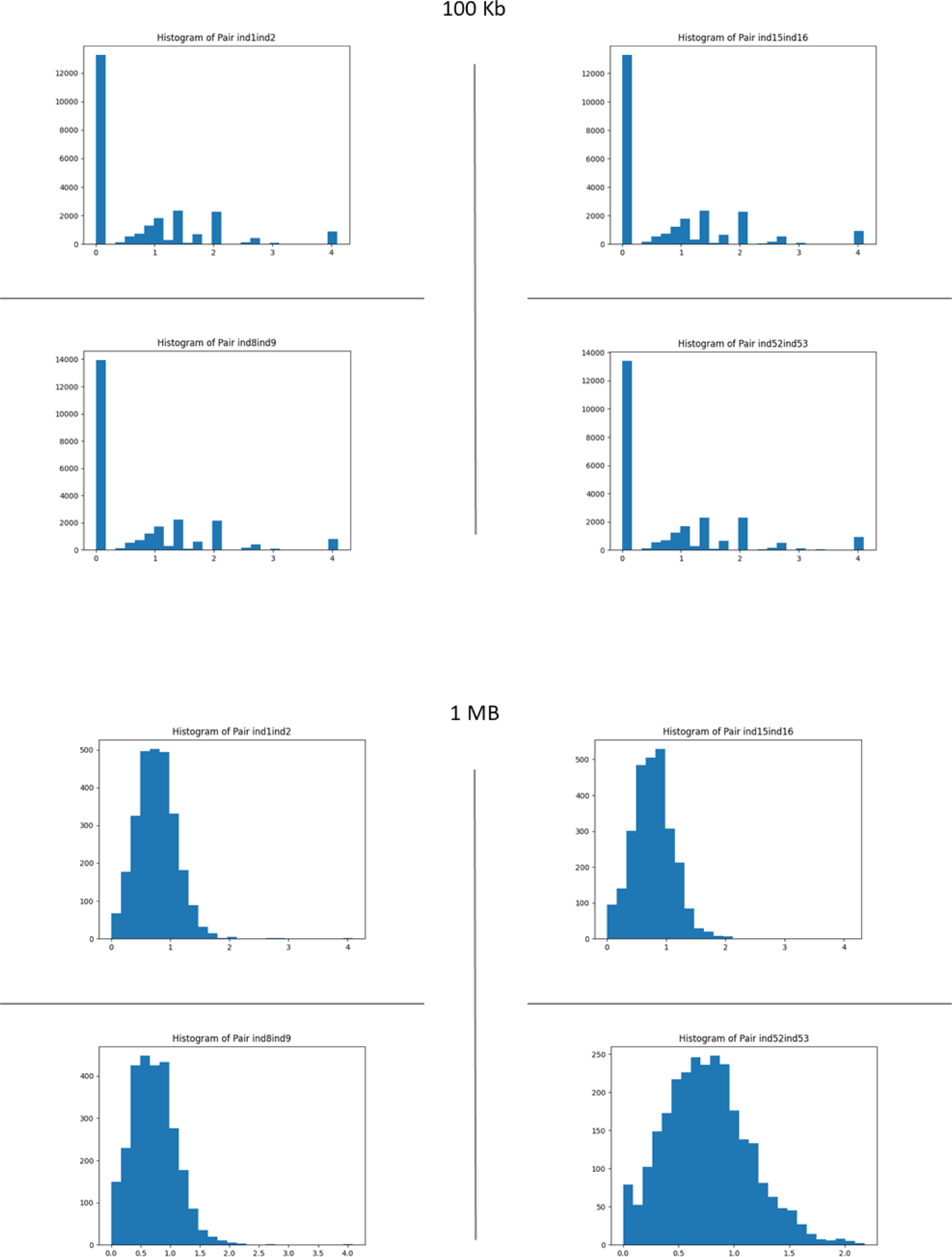
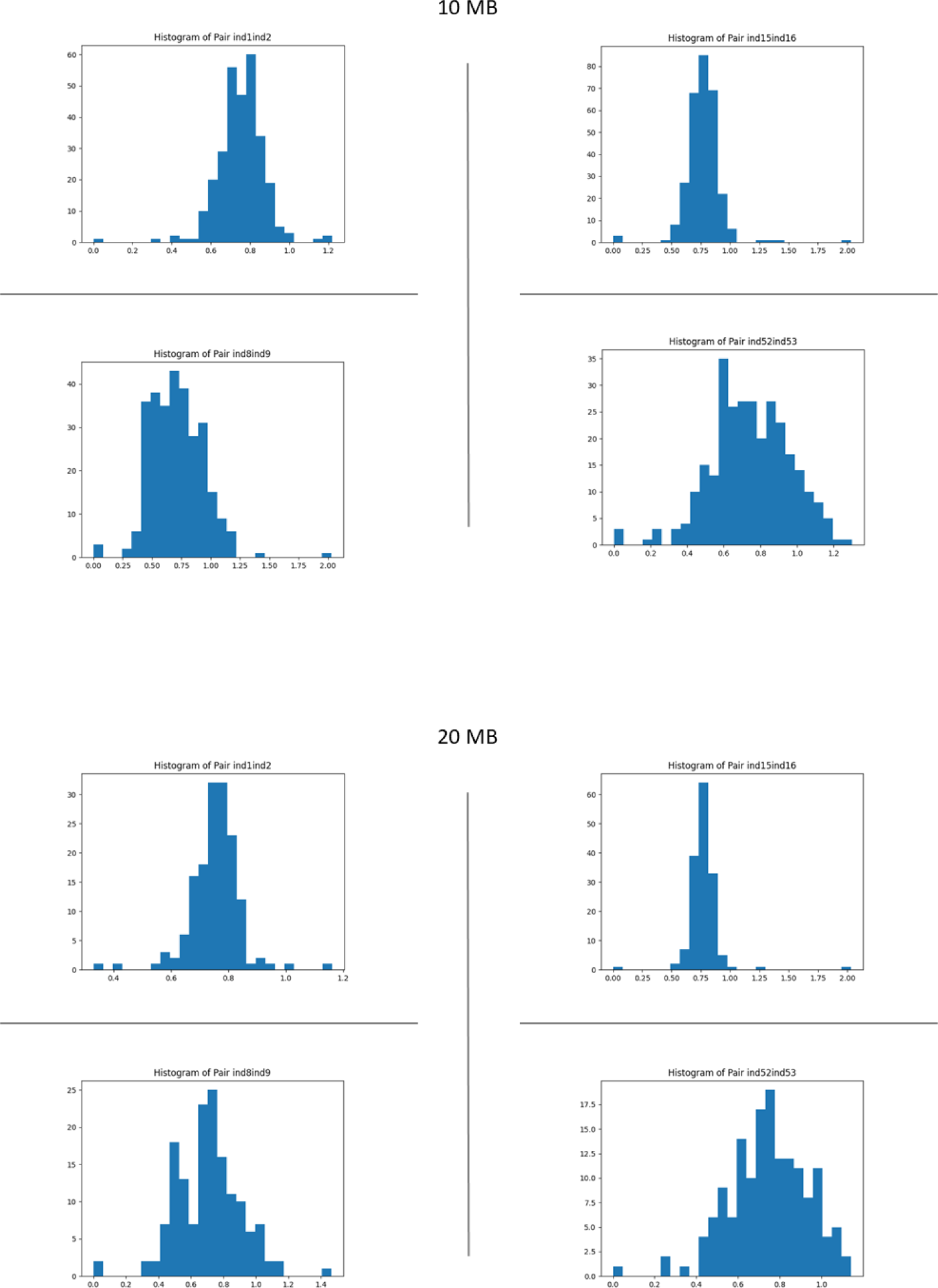
Examples of histograms of normalized P0 values for simulated parent-offspring (top) and sibling (bottom) pairs for varying window sizes. Smaller window sizes show very noisy distributions for parent-offspring and sibling pairs. However, windows of 20Mb result in distributions centered in the first-degree region with wider distribution for sibling pairs.

**Table S1:** Comparison of various available kinship inferring methods in terms of minimum coverage and minimum inferred relationship degree. The values shown here are retrieved from the original papers. We need to note that the numbers of SNPs and coverages are not always straightforward to compare between studies due to differences in (the size of) the underlying SNP panel and the background relatedness of the population.

| Method | Min. Coverage | Size of SNP panel used | Degree up to | Within Degree Differentiation | Type of Data Needed |
| --- | --- | --- | --- | --- | --- |
| IcMLkin [29] | 2X | 100,000 | Third | First degree (parent-offspring, siblings)<sup>‡</sup> | Biallelic genotypes, genotype likelihoods and population allele frequencies* |
| ngsRelate [30] | 1X | 100,000 | Third | First degree, some second degree (half-siblings) and some third degree (first cousins)<sup>‡</sup> | Genotype likelihoods and population allele frequencies** |
| TKGWV2 [31] | 0.026X | ~22,000,000 | Second | - | Pseudohaploid genotypes and population allele frequencies |
| KIN [32] | 0.05X | Not-specified | Third | First degree | Aligned reads*** (.bam files) |
| BREADR [33] | 0.04X | ~29,000,000 | Second | - | Pseudohaploid genotypes |
| ancIBD [35] | 0.25X (WGS) or 1X (1240k SNP capture) | ~1,200,000 | Sixth | First and second degree | Imputed genotypes**** |
| correctKin [34] | ~85,000 overlapping markers<sup>†</sup> | ~1,200,000 | Fourth | - | Pseudohaploid genotypes |
| READv1 [27] | 0.1X | ~1,200,000 | Second | - | Pseudohaploid genotypes |
| READv2 (this study) | 0.05X or ~500 SNPs (first degree)<br>0.1X or ~2,000 SNPs (second degree)<br>0.3X or ~15,000 SNPs (third degree) | 200,000 | Third | First degree | Pseudohaploid genotypes |

\*Estimated from the input data
\*\*Optional, can be estimated from the data
\*\*\*The required input files for KIN (the number of overlapping sites, the number of pairwise differences, and the probability of runs of homozygosity (ROH) in the windows where individuals have the same long allele sequence) are created by KINgaroo, which is provided together with the KIN package, by using these .bam files.
\*\*\*\*Imputed genotypes are created with GLIMPSE [64] by using aligned sequence data (.bam files) as a part of the pipeline.
<sup>†</sup> The authors of correctKin report their results with the term “overlapping markers”, which stands for the percentage of markers shared by both genotypes.
<sup>‡</sup> This classification is not automatically performed by the tool but can be achieved by comparing different statistics against each other.

**Table S2:** The number of Parent-Offspring and Sibling pairs present in the genotyping data in populations from the 1000 Genomes Project [40].

| Population Name | Number of Parent-Offspring Pairs | Number of Sib. Pairs |
| --- | --- | --- |
| ASW | 46 | 8 |
| CDX | 2 | NA |
| CEU | 1 | NA |
| CHS | 105 | 8 |
| CLM | 69 | NA |
| GBR | 2 | 1 |
| GIH | NA | NA |
| IBS | 100 | NA |
| KHV | 41 | 1 |
| LWK | 5 | NA |
| MXL | 60 | 3 |
| PEL | 70 | 1 |
| PUR | 66 | NA |
| TSI | 1 | NA |
| YRI | 112 | 4 |

**Supplementary Data 1** - READv2 results for the Rivollat data [22]. Explanations of the columns can be found in the second tab. This table also represents an example of the output of READv2.

